# Portable Arbitrary Pulse Generator for Driving μCoils for Micromagnetic Neurostimulation

**DOI:** 10.1101/2023.06.19.545512

**Authors:** Robert P. Bloom, Renata Saha, Zachary Sanger, Onri J. Benally, Kai Wu, Arturo di Girolamo, Walter C. Low, Theoden I. Netoff, Jian-Ping Wang

## Abstract

Micromagnetic stimulation (μMS) is a promising branch of neurostimulation technologies. Microcoil (μcoil) based magnetic stimulation uses micrometer sized coils that generate a time-varying magnetic field which as per Faraday’s Laws of Electromagnetic Induction induces an electric field on a conductive surface. This method of stimulation has the advantage of not requiring electrical contact with tissue, however these μcoils are not easy to operate. Large currents are required to generate the required magnetic field. These currents are too large for standard test equipment to provide, and additional power amplifiers are needed. To aid in the development and application of micromagnetic stimulation devices, we have created a compact single unit test setup for driving these devices called the μCoil Driver. This unit is designed to drive small inductive loads up to ±8 V at 5 A and 10 kHz.

## I. Introduction

In the field of neurostimulation, micromagnetic stimulation (μMS) is a promising alternative to electrical stimulation. Possible applications include the treatment of Parkinson’s, epilepsy, dystonia, tremors, and OCD [1, 2, 3, 4, 5, 6, 7]. Both methods stimulate neurons by generating an electric field, however they differ in how the electric field is created. Electrical stimulation generates electric field by applying a voltage across implanted electrodes, whereas μMS uses a time varying magnetic field to induce an electric field upon the neuron. Although electrical stimulation has provided desirable results, those electrical electrodes require galvanic contact with tissues leading to biofouling and encapsulation of the electrode [1]. Micromagnetic stimulation is an alternative that overcomes this drawback associated with electrodes, as they do not require galvanic contact with the tissue.

The devices used in μMS are popularly known as microcoils (μcoils). They are small coils of wire that generate a directed magnetic field when current flows through them. Custom coils can be fabricated as in Ref [8, 9, 10], or commercially available inductors as Ref [11, 12, 13], used in circuit designs, have been used [1]. In general, these coils tend to show very low impedances. This characteristic, along with the large currents required to produce suitable magnetic fields from the μcoil, make them difficult to drive. Standard test equipment cannot provide the required power and a power amplifier is required. A typical electrical setup for micromagnetic stimulation consists of a waveform source (function generator or digital to analog converter), power amplifier and occasionally a PC [11, 14, 13, 15]. Multi-unit test setups make transport more difficult during transition to clinical settings as they require more lab space.

Our goal was to condense the typical test setup into a single compact unit. A unified system allows for ease of testing and portability and is well suited for testing across multiple labs and enhances the ease of transitioning the technology into clinical settings. In addition, the unit is battery powered to help mitigate any interference or noise from the power lines. This is of significant advantage while recording electrophysiological signals from the brain as the recorded signals will not be affected much by power line noise, thereby facilitating ease of data analysis. In this work we present a portable, battery powered, 40 W arbitrary pulse generator designed specifically to work with the Magnetic Pen (MagPen) [11] set of devices, as well as other micrometer sized inductors. The prototyped unit reported in this work, better named as the μcoil Driver, provides the basic functionality required by the typical μcoil experiment. Future work may include expanding the functionality of the unit in terms of increasing the output power, adding additional output channels and synchronization to external inputs.

A similar driving circuit was described in [20]. Their unit was designed to drive similar coils using higher frequency continuous sin waves. The primary difference is the types of waveforms generated; the μcoil Driver is designed to generate arbitrary waveform pulses at lower frequencies.

The rest of this article is organized as follows. The design specifications, circuit design, MagPen fabrication and experimental setup are detailed in Section II. Section III covers the device operation and the results of animal testing. Discussion and conclusion of the results are presented in Section IV and Section V respectively.

## II. Methods & Procedures

The MagPen system, as reported by Saha et al. [11] is a micrometer sized coil (μcoil) used to stimulate neurons. Current is driven through the coils to generate a localized magnetic field, which in turn generates an electric field in the neurons. These coils operate in two regimes, low frequency activation and high frequency suppression. In the low frequency regime, typical operation consists of driving a 1-2 A sinusoidal pulse at frequencies of 1-5 kHz. At these frequencies the μcoils behaves as a resistor with parasitic inductance. Therefore, there is no appreciable phase shift and can be treated as a low resistance load. In most experiments the μcoils are driven up to 5 A (rarely, 10 A as reported in [12]) to evoke a response [11].

### A. Physical structure of the μcoil driver

The μCoil Driver’s physical design was influenced by the goals of small form factor and desired battery life. The overall size of the unit was largely determined by the chosen batteries, with extra space allotted for the circuitry and wiring (Figure 1(d)). The front of the unit, shown in Figure 1(a), features a touch screen display, status LED, output and USB ports and the power button. For ease of use the user is given a touch interface and menu style navigation, where user inputs and operations are reinforced by a status LED and audible beeps. The status LED displays whether the output is ON or OFF, as well as any system errors. Error messages are conveyed by the color and blinking of the status LED, beeping and error messages on the screen. A custom cable connects the output port to the μcoil. The USB port allows for arbitrary waveforms to be loaded from a PC. The back of the unit (Figure 2(b)) contains battery fuses, the battery charging port, external power port and the power selector switch. The power selector switch allows for the unit to operate using its internal batteries or from an external power source. The unit measures 152 mm wide, 255 mm long and 110 mm tall, and weighs 4.7 kg. This compact form factor allows for the μcoil driver to be easily transported between labs and to require minimal lab space. The unit is also small enough to fit in medium shielded test setups.

**Figure 1:**
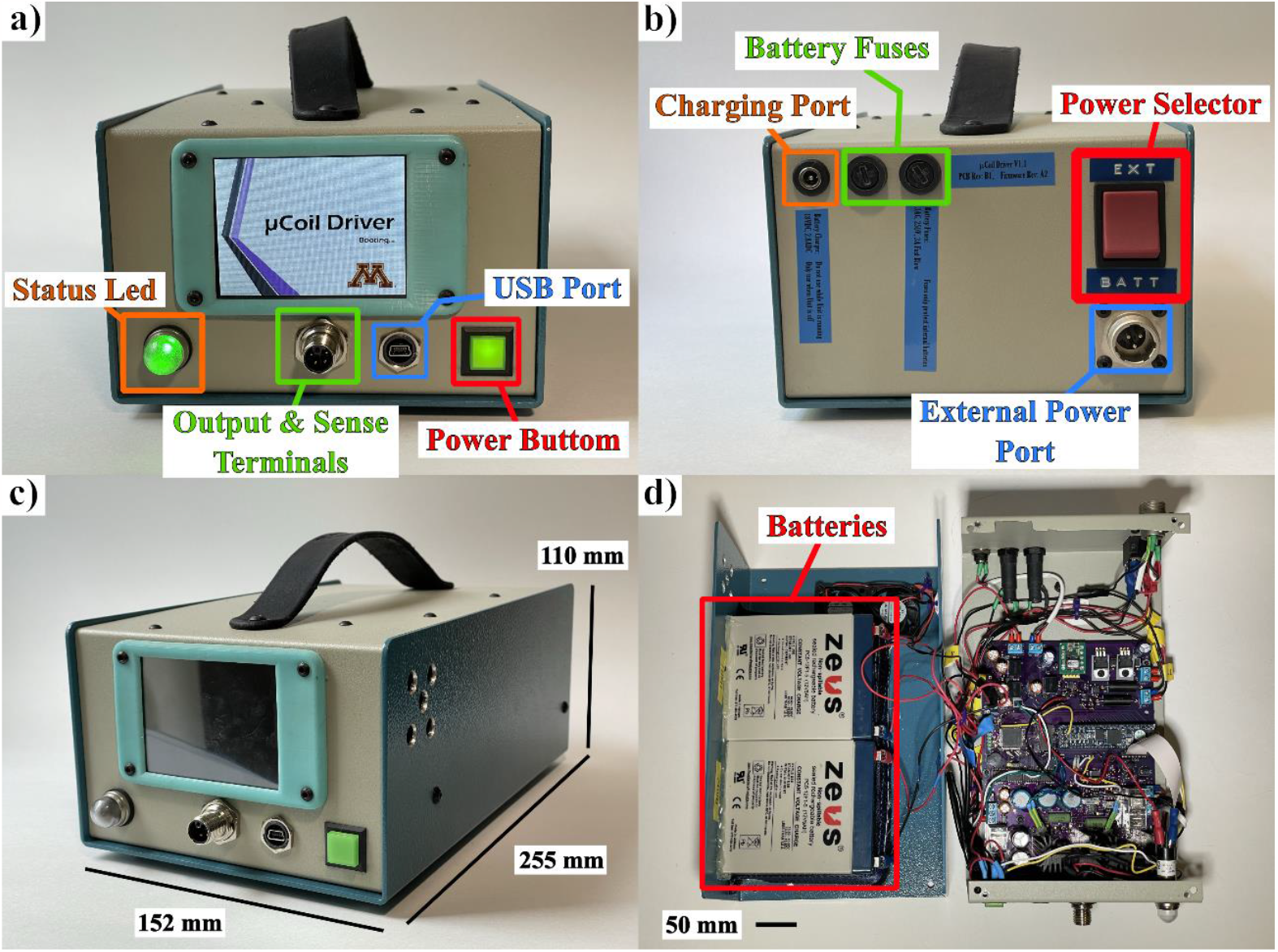
**a)** μCoil Driver front panel. **b)** μCoil Driver back panel. **c)** μCoil Driver unit. **d)** μCoil Driver internals.

**Figure 2:**
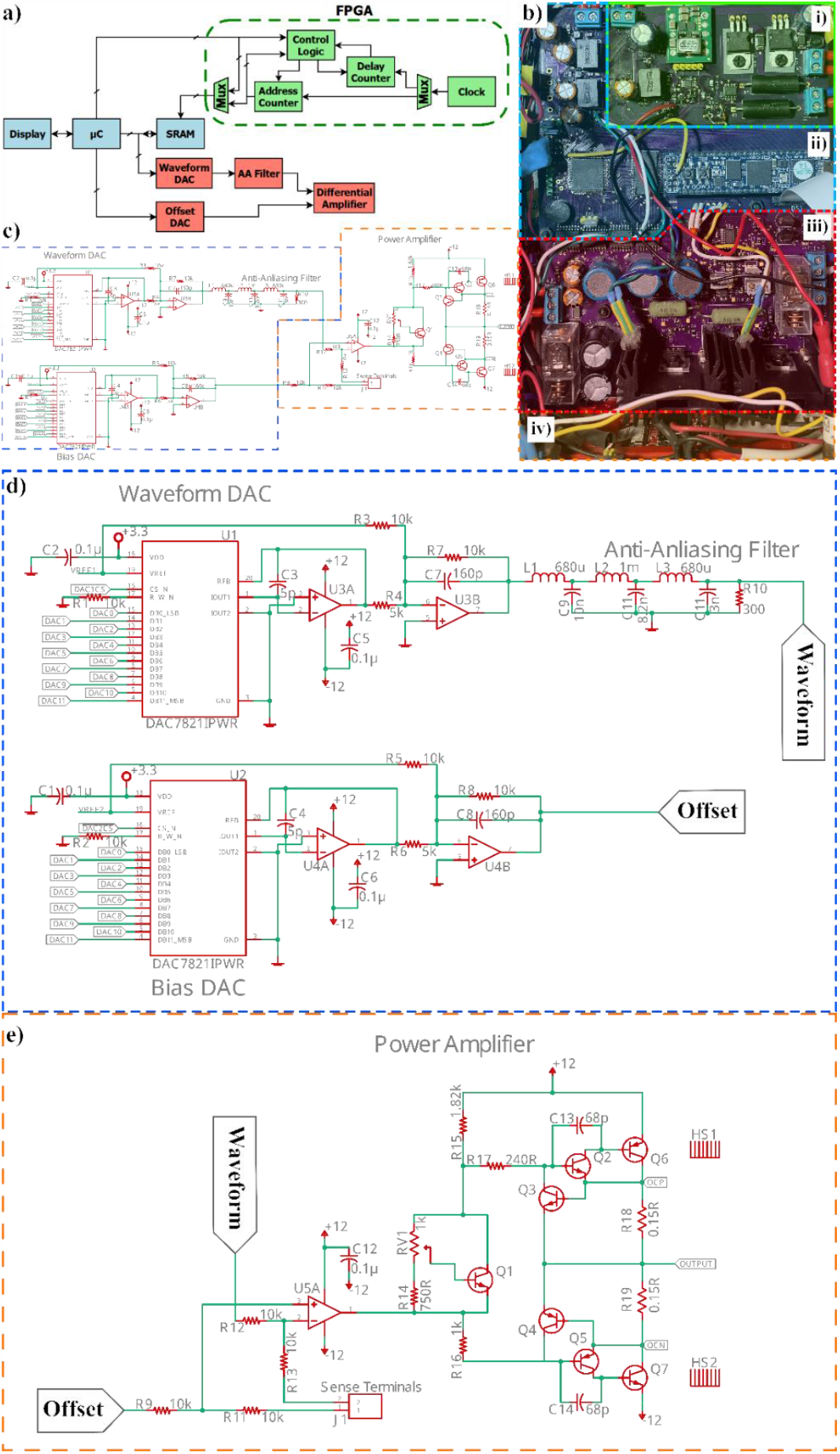
**a)** μCoil Driver functional diagram. **b)** PCB sections: **(i)** battery charger, **(ii)** digital section, **(iii)** analog section and **(iv)** display. **c)** Full analog signal path. **d)** Waveform generation section. **e)** Power amplifier section.

### B. μCoil Driver: Circuit Design

The μCoil Driver circuit can be broken into 3 main pieces (see Fig. 2(b)): battery charger, digital, and analog sections. The battery charger is a stand-alone circuit on the same PCB and is only active when the unit is in charging mode. The circuit charges the two 12 V batteries independently using a two-stage charging scheme. The charger is an analog circuit with a PIC12 microcontroller providing charge indication to the user and monitoring for even charging.

The digital section (see Fig. 2(ii)) of the μCoil Driver is responsible for running the graphical user interface (GUI), waveform calculation and system monitoring. These tasks are divided amongst three programmable devices: 4D Systems smart display, Xilinx Artix 7 FPGA and PSOC5 microcontroller. The 4D Systems display module is a touch screen LCD with integrated microprocessor. The interface was programmed using 4D Systems’ standard widgets and development software. It is responsible for running the GUI and passing information between the user and the PSOC5 microcontroller. The GUI allows for the user to select the output waveform and turn ON and OFF the output. The GUI also passes system information back to the user, like battery charge and error messages.

The user selected waveform is passed from the display to the microcontroller which then calculates and loads the waveform into a static random access memory (SRAM) IC. In remote load mode, the waveform is calculated by the PC and loaded onto the SRAM, where the microcontroller acts as an intermediary. The PSOC5 microcontroller is also responsible for monitoring the system. This includes managing the USB and UART communication interfaces, temperature and fan control, battery monitoring, managing various IC’s and output over current. To manage the large number of tasks and to allow for easy expansion, the PSOC5 microcontroller is running the FreeRTOS real time operating system [16]. The operating system runs with 1 ms ticks, and therefore precise timings and synchronizing the output is not possible.

Once the waveform is loaded into the SRAM the next step is to pass the data to the digital to analog converters (DAC). This action is completed by the FPGA, whose primary role is as a direct memory access controller (DMA). An FPGA was used for this ensure consistent timing. It also allows for future expansion to include synchronizing the output to an input trigger and to synchronize multiple units for driving arrays of coils. Once the data is transferred from the SRAM to the DAC’s the output waveform enters the analog section of the circuit.

The analog section of the circuit starts with the DAC and ends with the power amplifier. Signal path begins with the pair of DACs. One DAC is connected to the SRAM and is used for waveform generation. The second DAC generates a DC bias and is primarily used to remove any offsets that exist along the signal path. The output of the waveform DAC is passed through an anti-aliasing filter to remove the high frequency components of the quantized signal. The waveform and bias signals are combined using a differential amplifier. The differential amplifier was designed with a gain of 1 V/V and regulates the output voltage at the load using sense terminals. The differential amplifier consists of an operation amplifier (op amp) and a class AB power amplifier stage to increase the max output current. To help minimize any noise generated by the digital section, the analog and digital circuits have separate grounds and are connected via digital isolators IC’s.

The DAC’s operate at a sampling frequency of 200 kHz. The anti-aliasing filter is a 6th order, 50 kHz low pass Chebyshev filter. This high sampling frequency paired with the relatively low filter frequency produces a clean output with minimal distortion up to 10 kHz. To maximize the resolution through the full range of outputs, the DAC’s can switch between three references voltages: 2.048V, 4.096V and 8.192V. This allows the user to select the pulse’s amplitude with a resolution of 1 mV and eliminates the need for a variable gain amplifier.

For the power amplifier a class AB architecture was chosen to meet a balance between low distortion and power consumption. Sziklai pairs were used for the main driving transistors to maximize the output voltage swing and to increase the open loop gain of the differential amplifier. The differential amplifier is configured using sense terminals to regulate the voltage at the load. The use of sense terminals mitigates the voltage drop across long cables.

Over current events are detected by the sense resistors R50 and R51, and transistors Q8 and Q9. Sufficient current trough these resistors generates a voltage large enough to activate their corresponding transistors. When on, these transistors active to turn off the main drive transistors, thus clamping the output current. An identical set of transistors, placed in parallel (not shown in Figure 2), signal the microcontroller via an external interrupt. When an over current event is detected, the microcontroller removes power from the power amplifier stage.

### C. Fabrication of the MagPen prototype

The MagPen prototype was prepared for the demonstration of the uCoil Driver in the same way as that reported in Ref [11]. Inductors that were commercially available were soldered using solder flux and hot air blower on to the tip of a printed circuit board (PCB). The PCB thickness was thinned down to 0.4 mm into a semi-rigid structure, to facilitate an easy adjustment of the MagPen prototype over the nerve. Of the two kinds of MagPen prototypes reported in our earlier work [11], MagPen: Type V successfully activated the sciatic nerve. Just as reported earlier, a 10 μm thick Parylene-C coating was provided at the tip of the μcoil just to ensure the cause for sciatic nerve activation wasn’t from leakage current.

### D. Animal Surgery and Experimental set-up

Surgical methods and anesthesia regimens were conducted under a University of Minnesota Institutional Animal Care and Use Committee (IACUC) approved protocol 2004-38001A, similar to our work reported in [16]. Device testing was conducted in vivo in two Long-Evans (L/E) rats weighting 354.5 ± 94.05 grams. 1200 mg/kg Urethane was delivered through intraperitoneal injection and proper sedation assessed through toe pinch prior to local anesthesia subcutaneous delivery of 4 mg/kg Lidocaine and 2 mg/kg Bupivacaine at the site of incision. Urethane anesthesia was chosen over other approaches to preserve neurotransmitter release mechanisms. Heating devices were placed under the animal to maintain homeostasis while supplemental oxygen was administered via nose cone. A 30 mm incision perpendicular to the nerve exposed the dorsal-medial right quadricep muscular interstitial space. One cotton tipped applicator elevated the sciatic nerve from the interstitial space to space the nerve away from the muscle for effective placement of the μcoil. Throughout the surgery and micromagnetic coil stimulation, anesthetic depth and vital signs were checked every 15 minutes to ensure animal comfort and homeostasis.

## III. Results

### A. μcoil Driver: Operation

The μCoil Driver is powered by two 12 V batteries and can achieve maximum output swing of ±8 V, with a max peak current of 5 A. Under standard test settings of 1 kHz, 3 A pulses occurring every second, the unit is expected to have a battery life of 5 hrs. Alternatively, the unit can be easily switched to an external power source for longer duration tests.

The onboard GUI allows the user to select sinusoidal pulses. The pulse parameters are frequency, amplitude, phase, cycle count and interval (see Fig. 3(a) & (b)). Frequencies can be selected up to 50 kHz at a step size of 1 Hz. The amplitude (Vpeak) is selectable from 0 V to 8 V, in 1 mV steps. Phase can be chosen from 0° to 359° in 1° increments. Generated pulses idle at the last point. When the waveform is calculated, it will automatically offset the sinusoid such that it starts and stops at 0 V. For example, a sinusoidal pulse with a phase shift of 90° will generate a single sided negative pulse. Cycle count is the number of periods of the sinusoid per pulse and can be selected from 1-99 cycles (see Fig. 3(c)). Lastly, the interval is the delay between pulses. This can be selected from up to 10 sec at a resolution of 1 ms. Another feature is the ability to set the number of bursts. The user can select 1-5 bursts; each burst is customizable with the parameters above. When using multiple bursts (see Fig. 3(d)), each burst will be generated consecutively with the desired interval between bursts before repeating.

**Figure 3:**
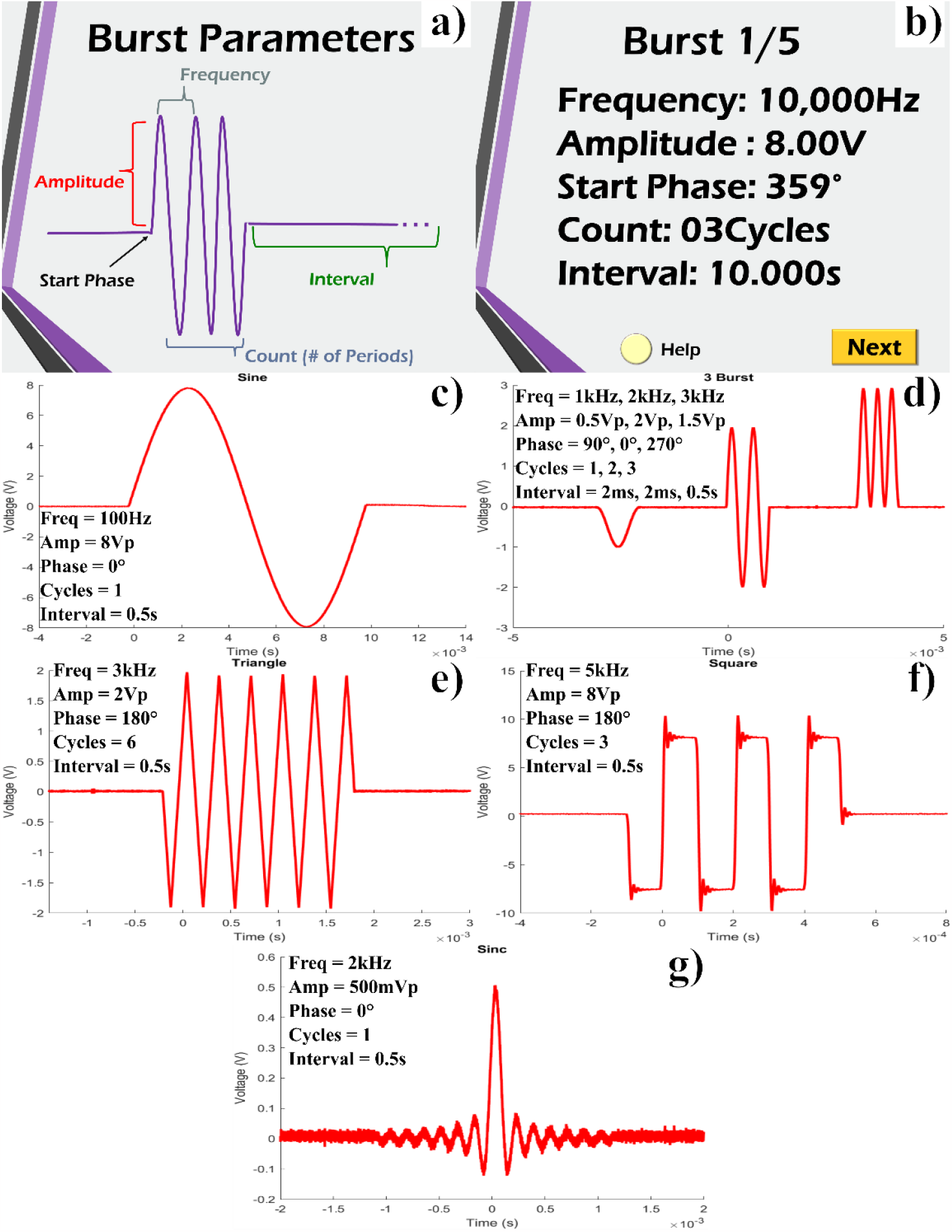
**a)** Waveform parameter definitions (help menu). **b)** Waveform selection menu. **c)** Sinusoidal output. **d)** Three burst outputs. **e-g)** arbitrary waveform outputs: triangle, square and sinc respectively.

Alternatively, arbitrary waveforms can be generated on a PC (see Fig. 2(e)-(g)) and passed to the unit via USB. The unit identifies to the PC as a COM (communication) port. MATLAB was used to generate the waveforms and load them onto the unit. The 8 Mb SRAM accommodates waveforms up to 2.5 sec in duration. Arbitrary waveforms must conform to the same limitations as the sinusoidal waveforms described above. Multiple bursts are also supported in this mode. The PC interface allows for basic control of the device such as turning output ON and OFF.

### B. μcoil Driver: Animal Testing

Testing consisted of validating the functionality of the unit, comparing to the test setup used by [11] for the MagPen system. The unit was validated by driving the MagPen devices to stimulate the sciatic nerve of a rat. Magnetic field pulses applied to the sciatic nerve can invoke a response from the rat. in the form of a sudden leg movement. Control tests were first done using bipolar electrodes (Plastics One Model No. MS303/8C used for electrical stimulation as well as the original MagPen driving system [11]. This gave a reference for the physical position of the MagPen device as well as the driving waveform parameters needed to invoke a response. Performing these control tests were extremely essential as there are several reports in the past portraying orientation dependent activation of neurons from these μcoils [11, 15, 12].

The complete experimental set-up with the portable μcoil driver powering the MagPen system held over the rat sciatic nerve has been shown in Fig. 4(a). Testing consisted of applying voltage pulses of varying wave shape and amplitude and looking for a response from the animal. The MagPen under test had a resistance of 1.8-2 Ω and an inductance of 0.6 μH. Pulse voltages were chosen to operate the μCoil Driver near its maximum current output of 3 A. Biphasic and monophasic waveforms were tested for the following waveforms: sinusoidal, square and triangular (see Table 1). Upon successful stimulation of the sciatic nerve, we see the limb muscle of the rat hind limb twitching. By ‘Pass’ (marked in GREEN) we mean a successful observation of the leg muscle movement. By ‘Fail’ (marked in RED) we mean that leg muscle movement was not observed upon MagPen stimulation when powered by the portable μcoil driver. Waveform parameters were set for a frequency of 1 kHz, count of 1 cycle, and interval of 1 sec. The amplitude of varied to show the sensitivity of the animal’s response. The tests were also repeated for phases of 0°, 90° and 270°. A phase of 0° represents a biphasic pulse whereas 90° and 270° are monophasic. The results of testing are shown in Table 1.

**Table 1:** Results of μCoil Driver and MagPen stimulation of rat sciatic nerve

| MagPen Sciatic Nerve Stimulation |  |  |  |  |
| --- | --- | --- | --- | --- |
| Frequency |  | Interval |  | Cycles |
| 1 kHz |  | 1 sec |  | 1 cycle |
| Sin |  | phase |  |  |
|  |  | 0° | 90° | 270° |
| amplitude | 5V<sub>pp</sub> | pass | fail | pass |
|  | 4V<sub>pp</sub> | pass | fail | pass |
|  | 2V<sub>pp</sub> | pass | fail | pass |
|  | 1V<sub>pp</sub> | fail | fail | fail |
| Triangle |  | phase |  |  |
|  |  | 0° | 90° | 270° |
| amplitude | 5V<sub>pp</sub> | pass | fail | pass |
|  | 4V<sub>pp</sub> | pass | fail | pass |
|  | 2V<sub>pp</sub> | pass | fail | fail |
|  | 1V<sub>pp</sub> | fail | fail | fail |
| Square |  | phase |  |  |
|  |  | 0° | 90° | 270° |
| amplitude | 5V<sub>pp</sub> | pass | pass | pass |
|  | 4V<sub>pp</sub> | pass | pass | pass |
|  | 2V<sub>pp</sub> | pass | fail | pass |
|  | 1V<sub>pp</sub> | fail | fail | fail |

**Figure 4:**
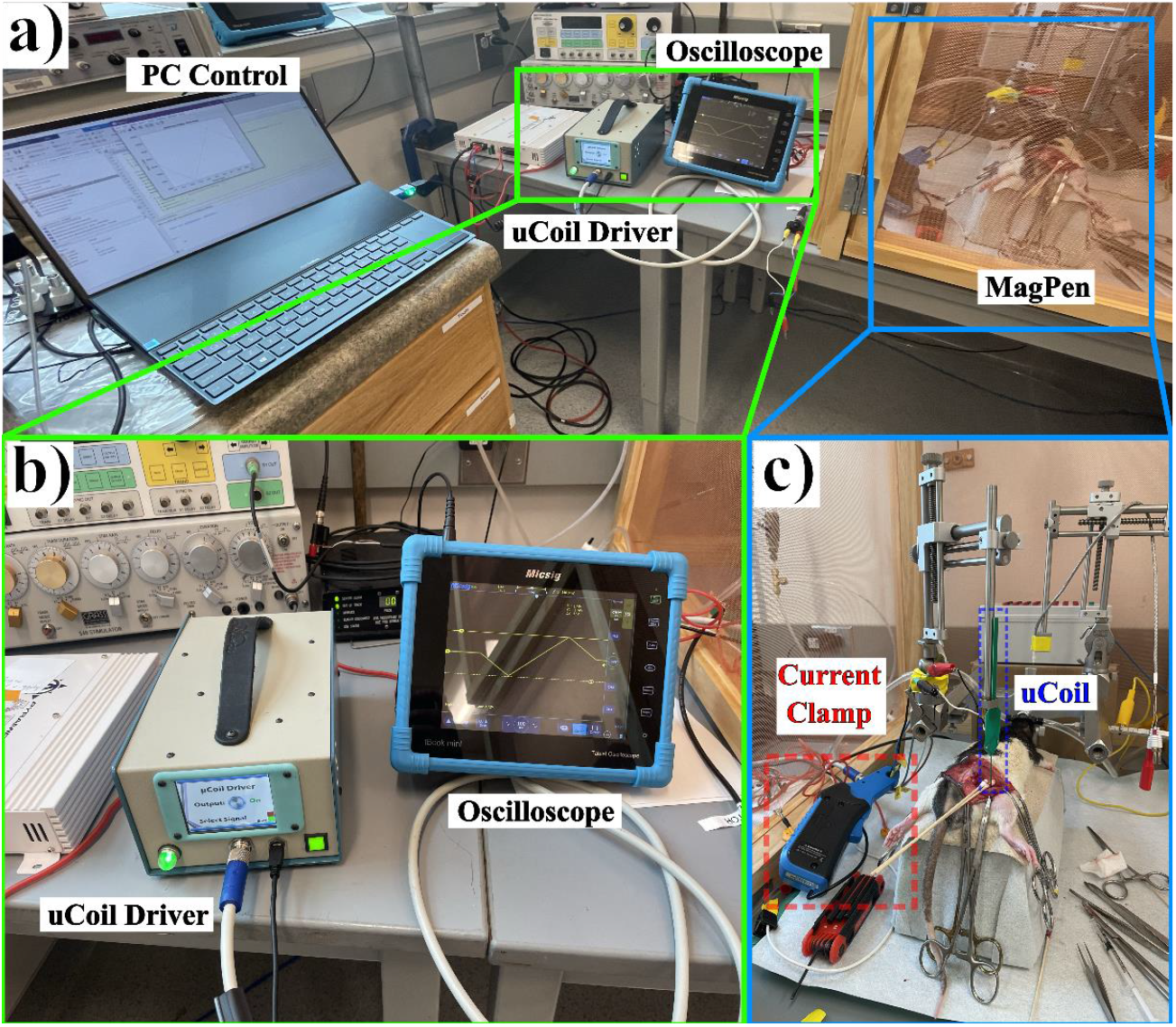
**a)** Experimental test setup consisting of PC, μCoil Driver, oscilloscope and MagPen **b)** μCoil Driver output on, with oscilloscope monitoring the output current. **c)** Surgery setup with MagPen μcoil placed over rat sciatic nerve.

From Table 1, we see that the MagPen stimulation of the sciatic nerve failed to evoke a leg muscle twitch for all waveform shapes and all phases of the 1 Vp-p amplitude driving the μcoil. This can be explained as follows: First, 1 Vp-p amplitude when driving a μcoil of resistance of about 2 Ω, means that only 0.5 A of current is powering the μcoil. This current is too low to activate the nerves. In contrast, 2 Vp-p, 4 Vp-p and 5 Vp-p when driving the μcoil, we observe leg muscle twitches at 0°and 270°but not at 90 °. This is because at 0°and 270 °, the waveforms start at 0 Vp-p and reach the positive maximum Vp-p at 1/4th of the total duration (here, total duration = 1/1 kHz = 1 msec; so, 1/4th of the total duration = 250 μsec); then, reach 0 Vp-p at ½ of the total duration (here, 500 μsec). Then again, reach the negative maximum Vp-p at 3/4th of the total duration (here, 750 μsec) until it finally reaches 0 Vp-p at 1 msec again. So, the nerve is expected to activate throughout the positive maximum and negative maximum during the durations of 500 μsec each. But, in case for the 90°phase (for sine and triangles), the waveform starts at positive maximum and then drops to 0 Vp-p at 250 μsec. In this short duration, the nerve cannot get activated, hence we do not see any leg muscle movement. For the square waves, we still observe leg muscle movement at higher amplitudes, 4 Vp-p and 5 Vp-p at 90°phase, because the square wave remains at the maximum positive and negative amplitudes throughout the 250 μsec durations and does not attenuate rapidly as for sine and triangular waves.

## IV. Discussion

Functionally, the μCoil Driver provides a more limited set of electrical outputs when compared to the original test setup. Its advantages are that it is a single unit, is battery powered, and has a small form factor. The onboard batteries are large enough to power the unit for up to five hours of continuous testing. For longer tests the unit can be switched to external power source. An additional advantage of being battery operated is that the unit can be isolated from noisy and intermittent wall power. The small size of the unit offers portability and can be used in small to medium shielded enclosures.

Another advantage comes from the power amplifier. The μCoil Driver uses a class AB amplifier whereas the original MagPen driving system utilized a class D amplifier [11]. The μCoil Driver can produce cleaner signals, without the characteristic high frequency noise of class D amplifiers. This design decision favored higher signal fidelity at the cost of lower driving power and shorter battery life.

Though the electrical characteristics of the μCoil Driver offer little benefit over the current setup, the μCoil Driver provides a basic platform which can be expanded upon. Future work may include higher output power, more channels, or synchronization between multiple units.

## IV. Conclusion

In this work we have presented the μCoil Driver; a portable, battery powered, 40 W arbitrary pulse generator designed specifically to work with the micromagnetic neurostimulators, facilitate the ease of carrying the driving unit across different experimental and clinical settings. The μCoil Driver can produce output waveforms (sine, square, triangular, arbitrary) with an upper limit of the voltage in the orders of ±8 V and a current and frequency rating of 5 A and 10 kHz. The unit is compact and easy to transport between labs. In vivo experiments on rat sciatic nerve stimulation further validated and experimentally demonstrated the μCoil Driver can effectively drive the MagPen system as efficiently as the standard test setup. This unit provides the basic functionality needed for micromagnetic stimulation operations and can serve as a basic building block on which other features can be added.

## Supplementary Video

**SV1:** Sine Wave, 1kHz, 10Vpp

**SV2:** Half Sine Wave, 1kHz, 10Vpp

**SV3:** Square Wave, 1kHz, 10Vpp

**SV4:** Half Square Wave, 1kHz, 10Vpp

**SV5:** Triangle Wave, 1kHz, 10Vpp

**SV6:** Half Triangle Wave, 1kHz, 10Vpp

**SV7:** Arbitrary Pulse, 2kHz, 12Vpp

## Notes

The authors declare no conflict of interest.

## Acknowledgements

This study was financially supported by the Minnesota Partnership for Biotechnology and Medical Genomics under award number ML2020. Chap 64. Art I, Sec11on 4. R.S. acknowledge the 3-year College of Science and Engineering (CSE) Fellowship awarded by University of Minnesota, Twin Cities. Research reported in this publication was supported by the University of Minnesota’s MnDRIVE (Minnesota’s Discovery, Research and Innovation Economy) initiative. The authors would also like to thank useful discussions Dr. Winfried A. Raabe, M.D. from the Department of Neurology; Kendall H. Lee, M.D., PhD, Charles D. Blaha, PhD and Yoonbae Oh, PhD from Mayo Clinic, Rochester, MN. Portions of this work were conducted in the Minnesota Nano Center (MNC), which is supported by the National Science Foundation through the National Nano Coordinated Infrastructure Network (NNCI) under Award Number ECCS-2025124. J.P.W, R.P.B. and R.S. also thank the Robert Hartmann Endowed Chair support.

